# A deep residual model for characterization of 5D spatiotemporal network dynamics reveals widespread spatiodynamic changes in schizophrenia

**DOI:** 10.1101/2022.11.16.516836

**Authors:** Behnam Kazemivash, Theo GM VanErp, Peter Kochunov, Vince D. Calhoun

## Abstract

Schizophrenia is a severe brain disorder with serious symptoms including delusions, disorganized speech, and hallucinations that can have a long-term detrimental impact on different aspects of a patient’s life. It is still unclear what the main cause of schizophrenia is, but a combination of altered brain connectivity and structure may play a role. Neuroimaging data has been useful in characterizing schizophrenia, but there has been very little work focused on voxel-wise changes in multiple brain networks over time, despite evidence that functional networks exhibit complex spatiotemporal changes over time within individual subjects. Recent studies have primarily focused on static (average) features of functional data or on temporal variations between fixed networks; however, such approaches are not able to capture multiple overlapping networks which change at the voxel level.

In this work, we employ a deep residual convolutional neural network (CNN) model to extract 53 different spatiotemporal networks each of which captures dynamism within various domains including subcortical, cerebellar, visual, sensori-motor, auditory, cognitive control, and default mode. We apply this approach to study spatiotemporal brain dynamism at the voxel level within multiple functional networks extracted from a large functional magnetic resonance imaging (fMRI) dataset of individuals with schizophrenia (N=708) and controls (N=510). Our analysis reveals widespread group level differences across multiple networks and spatiotemporal features including voxel-wise variability, magnitude, and temporal functional network connectivity in widespread regions expected to be impacted by the disorder. We compare with typical average spatial amplitude and show highly structured and neuroanatomically relevant results are missed if one does not consider the voxel-wise spatial dynamics. Importantly, our approach can summarize static, temporal dynamic, spatial dynamic, and spatiotemporal dynamics features, thus proving a powerful approach to unify and compare these various perspectives.

In sum, we show the proposed approach highlights the importance of accounting for both temporal and spatial dynamism in whole brain neuroimaging data generally, shows a high-level of sensitivity to schizophrenia highlighting global but spatially unique dynamics showing group differences, and may be especially important in studies focused on the development of brain-based biomarkers.

## 1 Introduction

Brain disorders affect many people worldwide every year. Schizophrenia is a chronic neuropsychiatric disorder, affecting over 20 million people worldwide. Schizophrenia often leads to cognitive and functional impairments and symptoms which typically emerge in late adolescence to early adulthood. Moreover, it shows a relapsing disease course in roughly seventy percent of the cases, so an early and rigorous diagnostic is critical to have a better treatment and get a reasonable clinical outcome [1, 2]. We still do not have understanding of the cause of schizophrenia, nor do we fully understand its impact on the brain and as a brain disorder. There is great interest in studying the underlying neural mechanism of psychosis. There have been numerous neuroimaging studies of schizophrenia [3, 4, 5, 6] in which the identified variations are highly complicated and distributed across many brain regions. However, existing, neuroimaging models of functional brain networks typically make strong assumptions about the associated variations in brain function. For example, most studies still do not allow for the possibility of time-varying changes in brain networks at the voxel level, i.e., spatial dynamics. Consequently, there are also almost no studies on the role of spatiotemporal brain dynamism effects in brain disorder as most are focused on static summaries or time-resolved variation in coupling among fixed nodes [7]. Generally, spatial brain dynamics refers to any changes in size, shape, or translation of active region over time, temporal dynamics refers to transient changes in coupling fixed brain regions over time, and spatiotemporal dynamics refers to transient changes in both the node/region and in its coupling to other nodes/regions [8]. Prior work focused on temporal dynamics has shown hypoconnectivity or dysconnectivity in transient coupling between functionally correlated sources for individuals with schizophrenia including transient changes in thalamic hyperconnectivity as well as hypoconnectivity between sensory networks and the putamen [9]. It has been shown that models that capture dynamics can improve sensitivity. For example, [10] showed incorporation of temporal dynamics improved classification accuracy for a three-way prediction of controls versus schizophrenia versus bipolar disorder. Other studies of temporal brain dynamism yielded promising results including 3 different brain networks exhibiting antagonism with severity of the illness. Despite the fact that schizophrenia is thought to involve complex morphological and functional dysconnectivity, there has been little work exploring potential spatiotemporal biomarkers that can distinguish patients from controls. One approach by [11] addressed the issue by utilizing a relatively constrained spatiotemporal model that achieved higher accuracy in comparison with classical methods. In this work, support vector machine classifiers were trained on functional connectivity dynamics and predicted patients versus controls with an accuracy of more than 90%. They also showed that constraining the model to ignore spatial or temporal dynamics yielded lower performance, with static functional connectivity having the lowest performing. Extensive research on voxel-wise spatiotemporal brain dynamism in schizophrenia is important to more fully characterize the underlying brain changes linked to schizophrenia. There are to date only a few studies that have begun to explore this. For example, prior work has evaluated the relevance of interactions between spatially distributed patterns or temporally synchronized brain networks described as spatiotemporal brain dynamics [12, 13]. Another study by [14] on saturated transient supra-network sources called polarization, showed remarkable differences in special patterns in time-resolved network connectivity such that high polarized sources are highly correlated with connection stability between auditory, sensory, motor, and visual networks. Other studies have begun to explore the relationship of spatial dynamics within a hierarchy of time-varying network components with different granularity levels where higher levels show more dynamism versus lower levels that illustrate more homogeneity [15]. Moreover, a study by [16], characterized the spatial chronnectome and highlighted cases where inter-network integration was changing over time, providing an important motivation to continue to extend such approaches for biomarker detection.

It is especially important to develop flexible approaches based on spatiotemporal brain dynamism to move towards a reliable biomarker for schizophrenia, but working on spatiotemporal dynamics is computationally intensive especially while utilizing deep learning models either in training or inference phase, but recent enhancements in computational infrastructures and algorithms like GPUs/TPUs and distributed systems [17] have made it possible to use deep learning techniques on fMRI data and study spatiotemporal brain dynamics. In this work we address issue by studying group differences using a novel framework with deep residual convolutional neural network models to estimate spatially flexible networks in 5D including space, time, and network. We apply this approach to a large study of schizophrenia patients and controls in order to evaluate the degree to which the 5D networks captured complex group differences.

## 2 Method and materials

We conducted our study by incorporating a framework called neuromark which leverages a fully automated spatially constrained ICA approach to estimate subject specific spatial maps and timecourses. In this work we used the neuromark_fMRI_1.0 template, consisting of 53 replicable brain networks to initialize a deep learning model which encodes spatiotemporal brain dynamism within an fMRI dataset to generate 4D voxel-wise dynamic brain networks each of the 53 networks, which are grouped into 7 domains including sensorimotor (SM), default mode (DM), auditory (AU), cognitive control (CC), visual (VI), subcortical (SC), and cerebellar (CB) as is shown in figure 1. This produces a 5D dataset including 53 4D brain networks, for each subject. Following this, we utilized different statistical metrics to summarize the network and to analyze group differences which are thoroughly discussed in the following sections.

**Figure 1.**
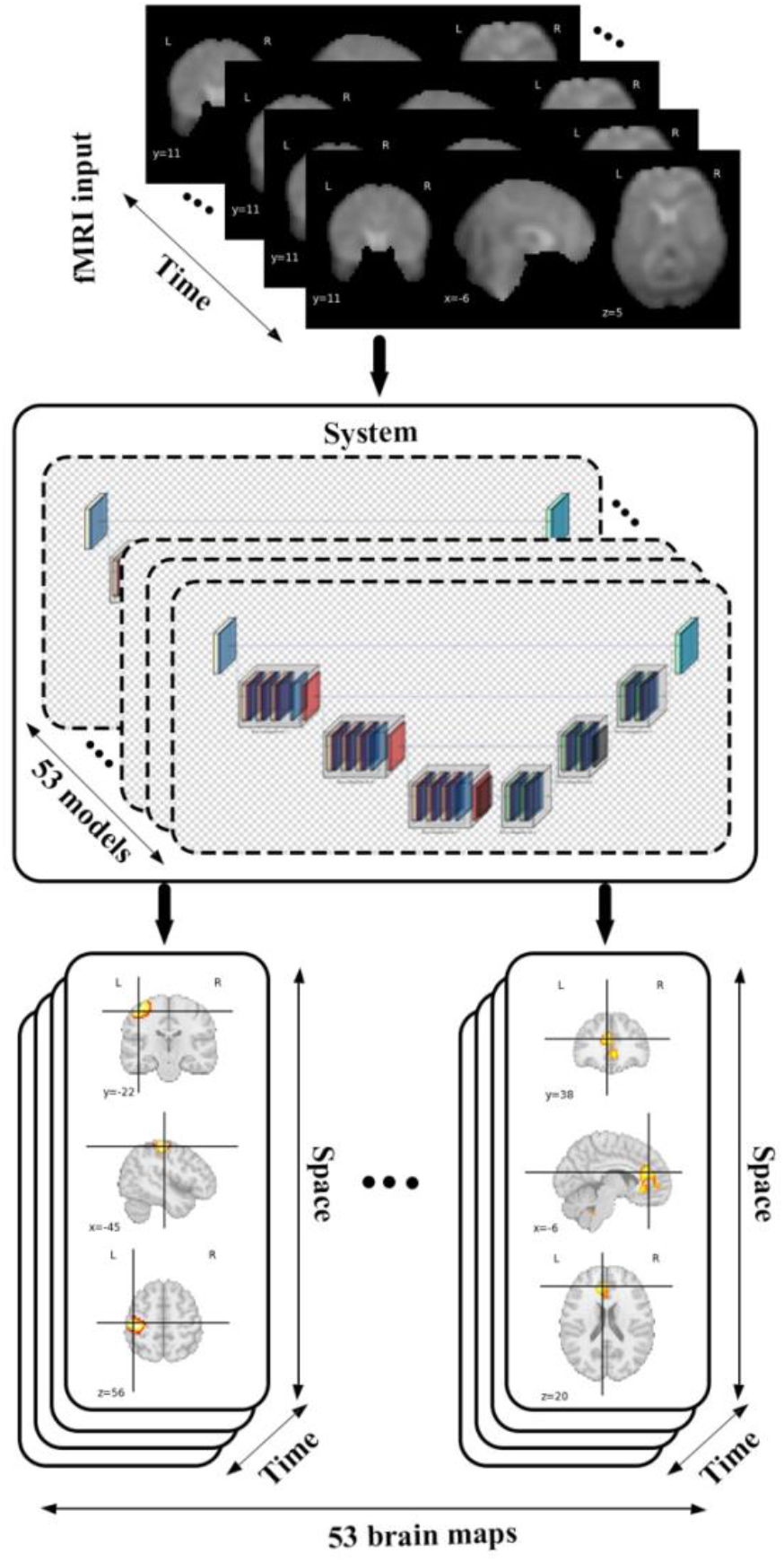
The diagram shows data flow in our framework by feeding an fMRI input into 53 different models each of which belongs to a specific brain domain like visual, cerebellar, auditory, sensory motor, default mode, cognitive control, subcortical leads to generating spatiotemporal maps (4D maps) for each of brain networks.

### 2.1 Demographic and image acquisition

We investigated potential differences between schizophrenia versus control groups by using 1218 resting-state functional magnetic resonance (fMRI) images from different existing datasets including 510 control and 708 schizophrenic subjects collected from two studies.

The first study utilized data collected at the MPRC (Maryland Psychiatric Research Center) project which was gathered by University of Maryland across three sites using 3-Tesla Siemens Allegra scanners with voxel size of 3.44 × 3.44 × 4 mm, field of view 220 × 220 mm, TR = 2000 ms, TE = 27 ms, and also 150 volumes; 3-Tesla Siemens Tim Trio scanner with voxel spacing size of 1.72 × 1.72 × 4 mm, field of view 220 × 220 mm, TR = 2000 ms, TE = 30 ms, and 444 volumes; and a 3-Tesla Siemens Trio scanner with voxel spacing size of 3.44 × 3.44 × 4 mm, field of view 220 × 220 mm, TR = 2210 ms, TE = 30 ms, and also 140 volumes. Also, similar subject inclusion criteria such as head motion less than 3° and spatial normalization, were applied to all participants [13, 18]. The second study was conducted on data collected on a multi-site study using 3T MRI scanners. During the resting fMRI acquisition, subjects were asked to relax and stay awake during the scan. Images were subjected to a harmonized preprocessing pipeline after excluding subjects with head motion greater than 3mm in x, y, and z or 3° in pitch, roll, or yaw. All participants signed an informed consent form based upon guidelines of the Internal Review Boards of corresponding institutions and expert psychiatrists diagnosed the schizophrenia patients. The demographic information of the collected data is shown in Table 1.

**Table 1.** Demographic Information of Data.

| Project | Diagnosis | Sex | Samples | Age (years) |  |
| --- | --- | --- | --- | --- | --- |
| | | | | Mean ± sd | range |
| Study 1 | CNT | M | 100 | $38.841 \pm 14.21$ | 12.0 - 77.0 |
| | | F | 154 | $41.0 \pm 16.31$ | 10.0 - 79.0 |
| | SCZ | M | 111 | $35.09 \pm 12.93$ | 13.0 - 63.0 |
| | | F | 52 | $43.05 \pm 13.64$ | 13.0 - 63.0 |
| Study 2 | CNT | M | 133 | $28.85 \pm 7.82$ | 18.16 – 45.75 |
| | | F | 123 | $28.40 \pm 7.82$ | 17.66 – 45.33 |
| | SCZ | M | 297 | $28.03 \pm 7.54$ | 16.33 – 46.75 |
| | | F | 248 | $30.28 \pm 7.66$ | 16.83 – 54.0 |

### 2.2 Data preprocessing

Data preprocessing plays a crucial role in neuroimaging analysis and can significantly impact the interpretation of result. In this work, standard preprocessing steps were applied to all resting state fMRI data utilizing statistical parametric mapping toolbox [19] including removal of the first 5 timepoints for magnetization equilibrium, head motion correction, and slice timing correction. Moreover, data were spatially normalized to the Montreal Neurological Institute (MNI) Echo-planar imaging (EPI) template, resampled to 3 × 3 × 3 mm, and smoothed by a 6mm Gaussian kernel (FWHM = 6 mm).

### 2.3 Model structure

Spatiotemporal brain dynamism forms the backbone of our group comparison study, characterized by a 5D set of brain networks extracted from fMRI data. We incorporated a brain parcellation framework [20, 21] including 53 pre-trained models each of which produces a score map (probabilistic map) for a specific brain network, varying over space, time and across subjects and consequently is able to encode spatiotemporal brain dynamics. Each of the models is a residual U-Net style regressor containing 36 layers such as 3D convolution, transposed 3D convolution, max pooling, batch normalization, and dropout layers grouped into encoding and decoding blocks and was trained and evaluated using 1470 samples (volumes) from a subset of 3 preprocessed fMRI images in the UK Biobank dataset as the input and relevant extracted ICA maps as priors due to supervised training approach. Furthermore, mean squared error (MSE) as the loss function, Adam optimizer with adaptive learning rate of 0.00001 and step size of 5 were utilized to train models in 200 epochs along with a volume-based data feeding policy and all volumes were normalized using min-max normalization method to get a faster convergence in training process. An early stopping method is also applied to have a better generalization and prevent the overfitting issue. We also fine-tuned the pre-trained model using 49,000 samples (volumes) from a subset of 100 preprocessed fMRI data in same dataset, and results were similar to that from the initial model.

The structure of the model was configured based on two main residual encoding and decoding blocks, wrapping different layers. In the proposed configuration, all encoding blocks have identical structure inside the block but different out-channel sizes between blocks. First encoding block has 3 volumetric convolution layers with same out-channel size between layers followed by a batch normalization layer after each of them. There is also a 3D max pooling layer with stride of 1 and kernel size of 3 after each encoding block and eventually a drop out layer with ratio of 0.5 as the last component of encoder segment. Besides, there are 3 decoding blocks in decoder segments and each of them contains a couple of 3D convolution transposed layer followed by a batch normalization layer. Schematic diagram of the model structure and layer details are given in figure 2 and table 2.

**Table 2.** Model architecture.

| Layer (type) | Output Shape | Param # | Tr.Param # |
| --- | --- | --- | --- |
| Conv3d-1 | [5, 64, 51, 61, 50] | 1792 | 1792 |
| Sigmoid-2 | [5, 64, 51, 61, 50] | 0 | 0 |
| ResEncBlocks-3 | [5, 32, 47, 57, 46] | 110880 | 110880 |
| ResEncBlocks-4 | [5, 16, 43, 53, 42] | 27792 | 27792 |
| ResEncBlocks-5 | [5, 8, 39, 49, 38] | 6984 | 6984 |
| Dropout3d-6 | [5, 8, 39, 49, 38] | 0 | 0 |
| ResDecBlocks-7 | [5, 16, 43, 53, 42] | 10464 | 10464 |
| ResDecBlocks-8 | [5, 32, 47, 57, 46] | 41664 | 41664 |
| Dropout3d-9 | [5, 32, 47, 57, 46] | 0 | 0 |
| ResDecBlocks-10 | [5, 64, 51, 61, 50] | 166272 | 166272 |
| ConvTranspose3d-11 | [5, 1, 53, 63, 52] | 1729 | 1729 |
| Sigmoid-12 | [5, 1, 53, 63, 52] | 0 | 0 |
| Total params: 367577 |  |  |  |
| Trainable params: 367577 |  |  |  |
| Non-trainable params: 0 |  |  |  |

**Figure 2.**
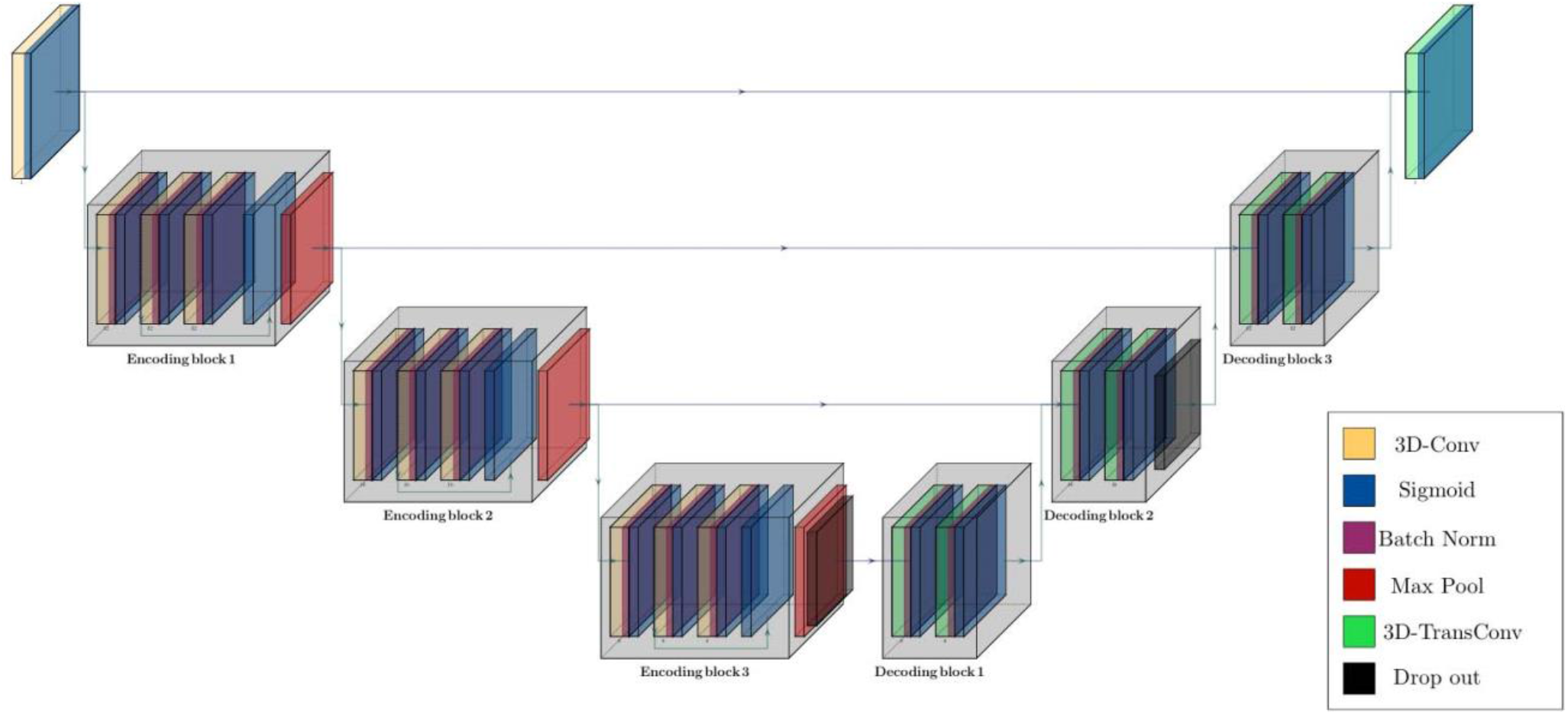
Schematic diagram of residual parcellation model with U-Net style structure including encoding and decoding blocks that contain combination of layers such as 3D convolution, max pooling, batch normalization, drop out, and convolution transposed. Black arrows show skip connections and data flow in and outside of each block.

### 2.4 Statistics and evaluation metrics

In this work we evaluate group differences between patient and control data considering multiple fully fluid 4D networks. Consequently, it is crucial to have solid and reliable metrics for interpreting brain dynamism differences between groups of subjects in both spatial and temporal aspects. More specifically, we study differences in spatial maps between control versus schizophrenia subjects by generating 4D maps for each of networks.

Furthermore, we analyze spatial variability over time by computing spatial deviation over time for each of control and schizophrenia groups on multiple summary measures. To do this, we calculate summation of absolute consecutive timepoints differences, and then averaging over subjects as is given in the following equations:

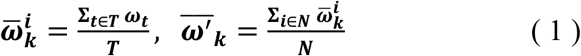

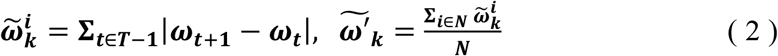

Here, 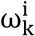 is the output of the model for brain network *k* and subject *i* with same shape as the input fMRI data (4D tensor), *T* denotes number of timepoints in ω. Also, 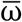shows voxel-wise average over time and, 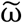denotes sum of absolute differences between consecutive timepoints. Eventually, 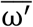and 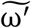 refer to averaged values over subjects and both functions result in a 3D tensor for each one due to voxel-wise operations. Moreover, we use a voxel-wise T-test for comparing our maps to identify regions showing group differences between controls versus patients using the following equation:

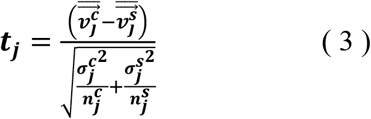

Where 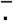shows average value, t_j_ refers to t-Test value of a specific voxel, 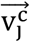is a vector containing different values of a specific voxel in 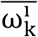or 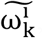, and also 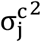is the variance of relevant vector for all controls. Also, 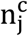denotes number of control subjects. We have same definition for 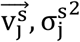and 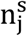in patients with schizophrenia and obtain a 3D tensor after computing *t* for all voxels.

We can also compute measures that capture temporal brain dynamism by evaluating the temporal coupling among networks using static functional network connectivity (sFNC) or dynamic version of that (dFNC) between each pair of spatiotemporal maps, ordered to show functional interactions across different brain networks. To do this, we compute the Pearson correlation between each pair of spatiotemporal maps using the whole length scan to get static functional network connectivity (sFNC) which eventually forms a matrix with size of k× k (number of all maps) for each subject by calculating sFNC for all pairs [22].

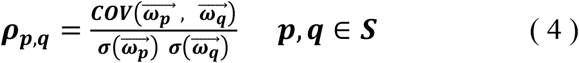

Here, *S* is a set of all extracted brain networks, 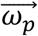and 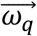are generated 4D maps for network *p* and *q* which are flattened into vectors and contain all brain voxels at all timepoints.

Recent studies emphasize on dynamic nature of functional connectivity in BOLD fMRI data in both animals and humans [4, 23, 24]. In line with this, we can compute dynamic functional network connectivity (dFNC) which highlights FNC variation over time, by applying same procedure as sFNC on a subset of timepoints with a constant window size that has overlap with the previous subset with ratio of α to estimate multiple FNC matrices for an individual. We can then use clustering to identify transient recurring patterns of functional connectivity, called functional states, and summarize these in various ways including the occupancy ratio (OR) which provides the time percentage of each state occurring during a scan [25].

## 3 Experimental results

We incorporated our BPARC framework with 53 pre-trained models to extract individual 4D spatiotemporal brain maps each of which belonging to one of sensory motor (SM), default mode (DMN), auditory (AU), cognitive control (CC), visual (VI), subcortical (SC), and cerebellar (CB) domains for all 1218 subjects. Then, we studied group level differences between control and schizophrenia subjects regarding to averaged spatial maps, averaged spatial dynamics, and eventually static and dynamic functional network connectivity (FNC).

### 3.1 Group differences in spatial maps

In the proposed study, we inspect group level dissimilarities in spatial maps between healthy-control and patients with schizophrenia by averaging maps over time and subjects, then computing 2-sample t-tests to capture group differences as shown in figure 3 for a sample subset of networks. The resulting maps depict plausible representation as we can see homogeneous parcels — connectivity in relevant brain regions for all averaged spatial maps as shown in figure 3.

**Figure 3.**
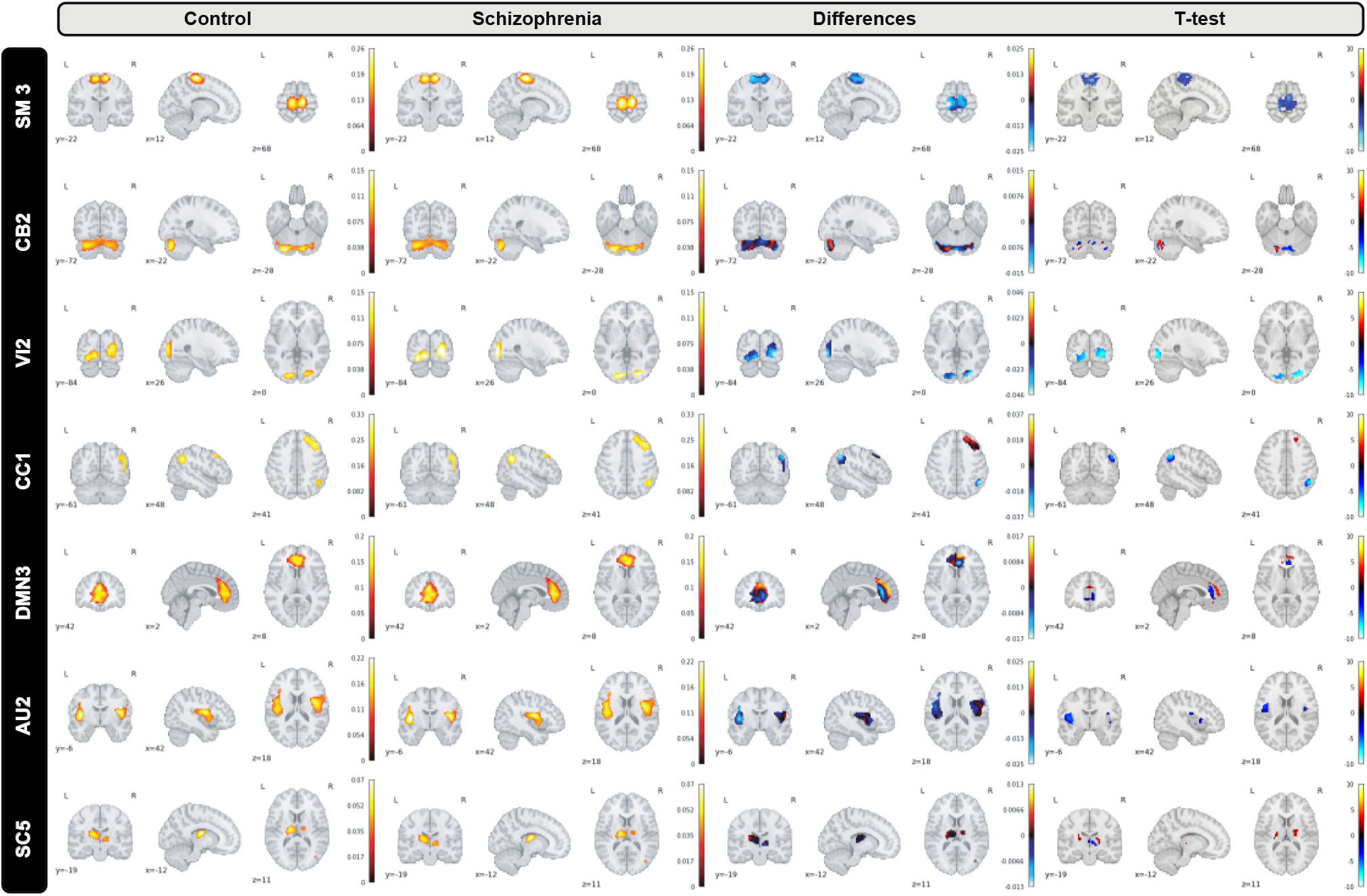
A subset of group level spatial maps averaged over time and subject for each control and schizophrenia patient along with subtracted maps and 2 sample T-tests to illustrate group level dissimilarities in sensori-motor (SM3), cerebellar (CB2), visual (VI2), cognitive control (CC1), default mode network (DMN3), auditory (AU2), and subcortical (SC5) networks. Furthermore, all brain maps are masked to focus on region of interest (ROI).

Moreover, we can see most networks show significant differences between the patients and controls, indicating widespread differences in brain connectivity between the two groups. For example, we see higher amplitude in voxels within sensori-motor-3 and visual-2 networks in schizophrenia patients. We also see other networks which show higher amplitude for the control group, for example in default mode network-3. Two-sample t values computed by applying 2 sample T-test highlight group differences across the various networks. In order to have a deeper insight into spatial maps in both groups, we carried out a statistical analysis on distribution of peak voxel within region of interest (ROI). Hence, we located 2 peak voxels in positive and negative segments of difference map and gathered values of those voxels in all subjects and drew a violin plot to see how distribution of those coefficients vary regarding median, min, max, and interquartile range (IQR) as shown in figure 4.

**Figure 4.**
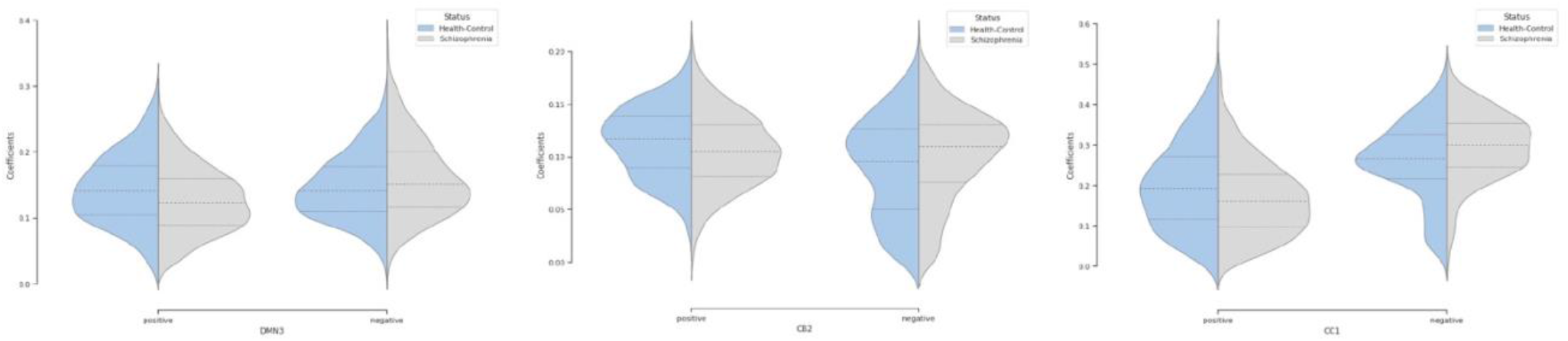
Violin plot shows distribution of peak voxels in positive and negative segments for all subjects in default mode network (DMN3), cerebellar (CB2), cognitive control (CC1) networks.

Furthermore, we inspected spatial dynamics for both control and schizophrenia groups. One summary measure for evaluating spatial dynamics is group level spatial deviation over time. Precisely, it can be defined as summation of absolute differences between consecutive timepoints, averaged over subjects and shows voxel-level activity patterns over time. Results show interesting information and patterns that is not captured by the overall mean activity (not visible in averaged maps) as is shown in figure 5 for a subset of brain networks. Interestingly, our observation illustrates functional connectivity near the boundaries of the active region in the cerebellar-2 network which is not visible in averaged maps or sensori-motor3 network in which we can see a transient linking to the cerebellum.

**Figure 5.**
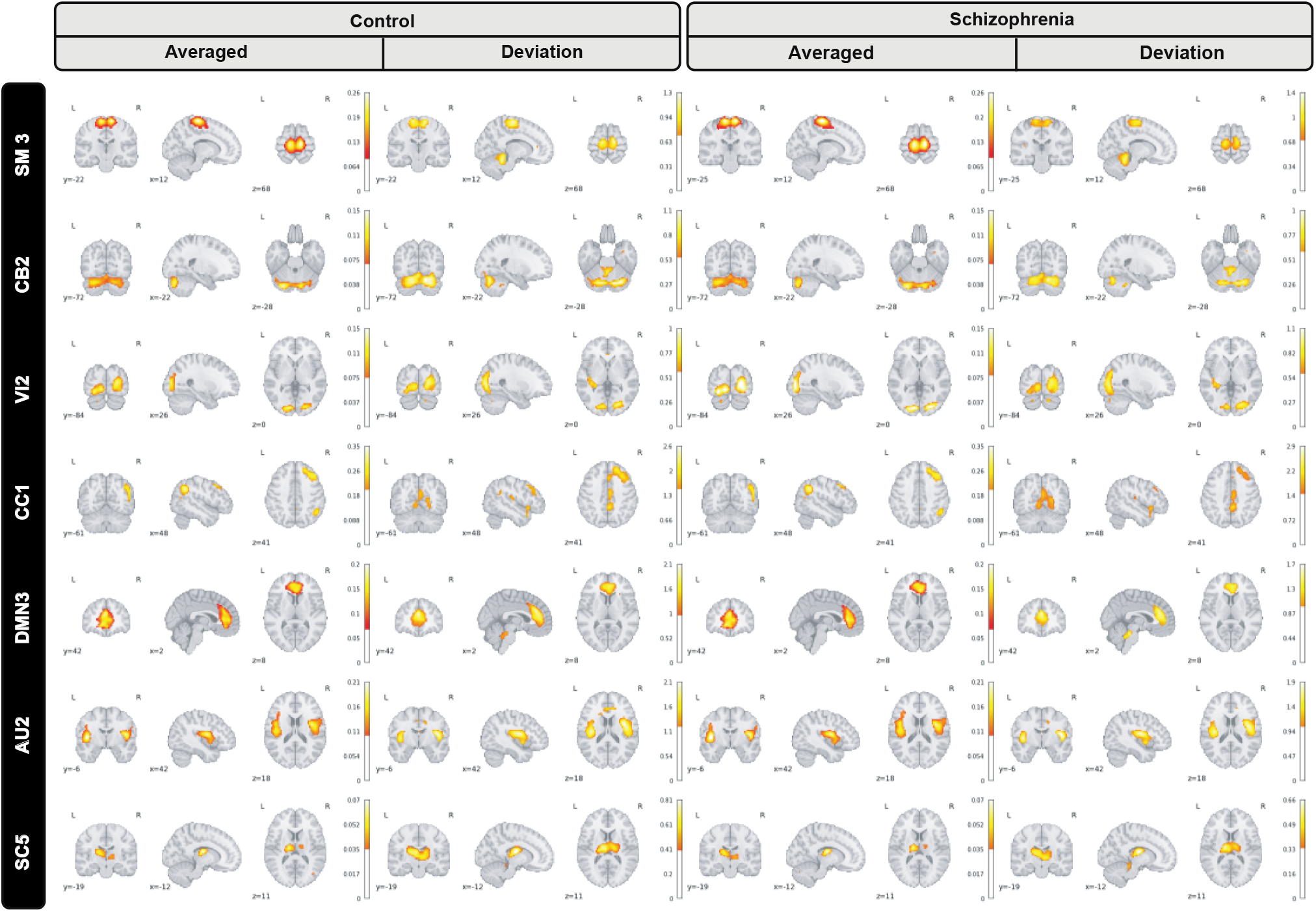
Group level spatial dynamics which change over time, averaged over subjects for the control schizophrenia in sensori-motor (SM3), cerebellar (CB2), visual (VI2), cognitive control (CC1), default mode network (DMN3), auditory (AU2), and subcortical (SC5) networks. All brain maps are thresholded to highlight functional changes which are not visible in the averaged maps.

### 3.2 Group differences in FNC

We also summarized the functional behavior of the 4D brain networks for both groups of healthy-control and schizophrenia via static functional network connectivity (static-FNC) which is Pearson correlation between all voxels in all timepoints for all 53 networks. We also show 2-sample t-test on the FNCs (thresholded at p<0.05 corrected for multiple comparisons via the false discovery rate) to highlight group level differences as is shown in figure 6.

**Figure 6.**
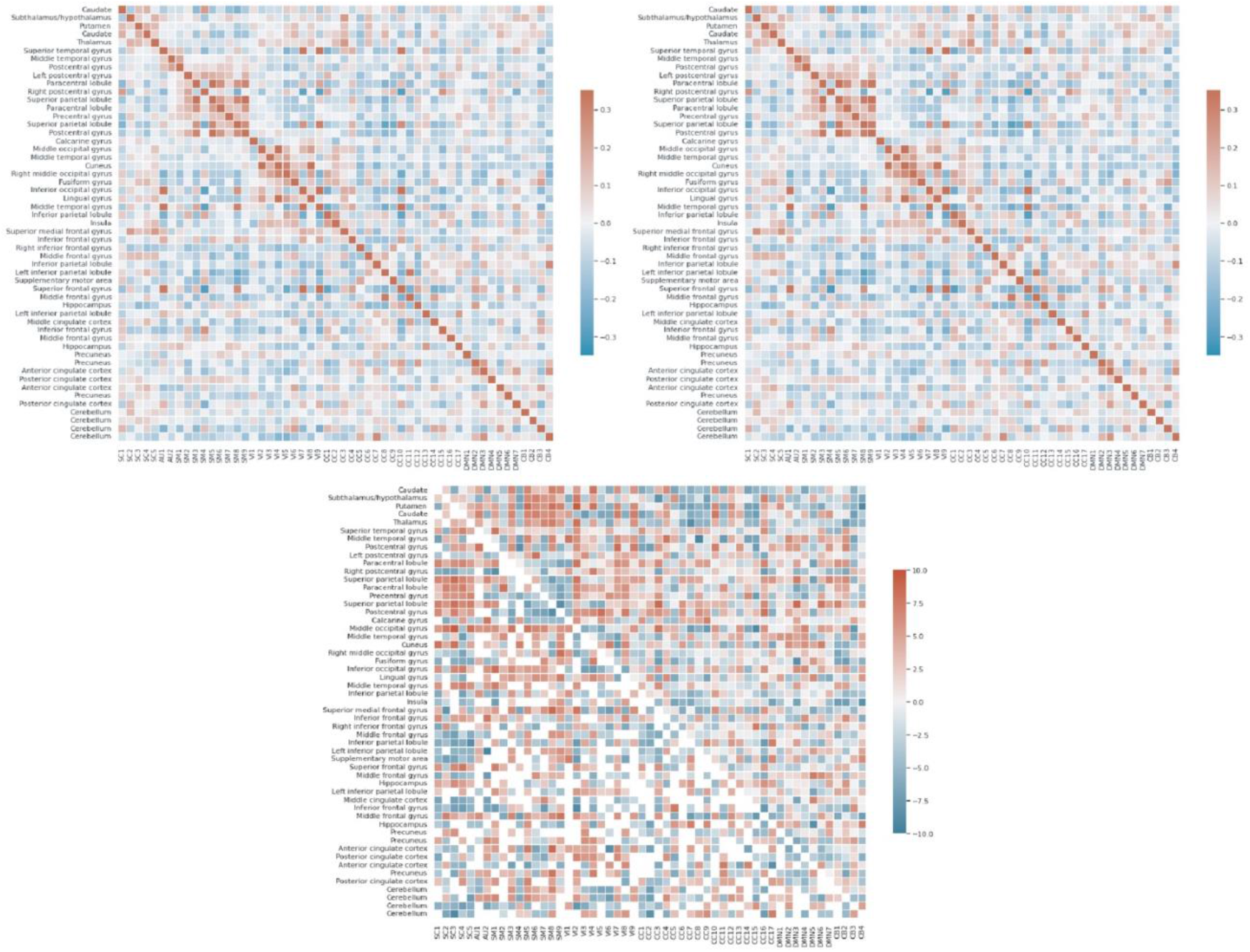
Each of FNCs illustrates correlation between all 53 components grouped into 7 sub-domains including default mode network (DMN), visual (VI), auditory (AU), cognitive control (CC), subcortical (SC), sensori-motor (SM), and cerebellar (CB) for controls (top left), schizophrenia patients (top right), and thresholded 2 sample T-test for HC-SZ (bottom).

In addition to static-FNC, we characterized the group level temporal coupling and dynamics by comparing the occupancy ratio in dynamic functional network connectivity between schizophrenia and healthy-control groups. To do this, we generated dynamic-FNCs by using windows size of 30 and overlapping ratio of 10 to generate various number of windows for each of subjects. Next, we applied k-means algorithm to cluster all generated windows into a set of separate clusters such that distance of each window within a cluster to the cluster centroid is minimized and the optimal number of clusters was estimated by elbow criterion. This procedure resulted in recognizing 4 initial states which are verified by centroids and used to predict label of all dynamic FNC windows for each of subjects. Finally, we computed the OR by counting frequency of states for each of subjects as is shown in figure 7 with all initial states.

**Figure 7.**
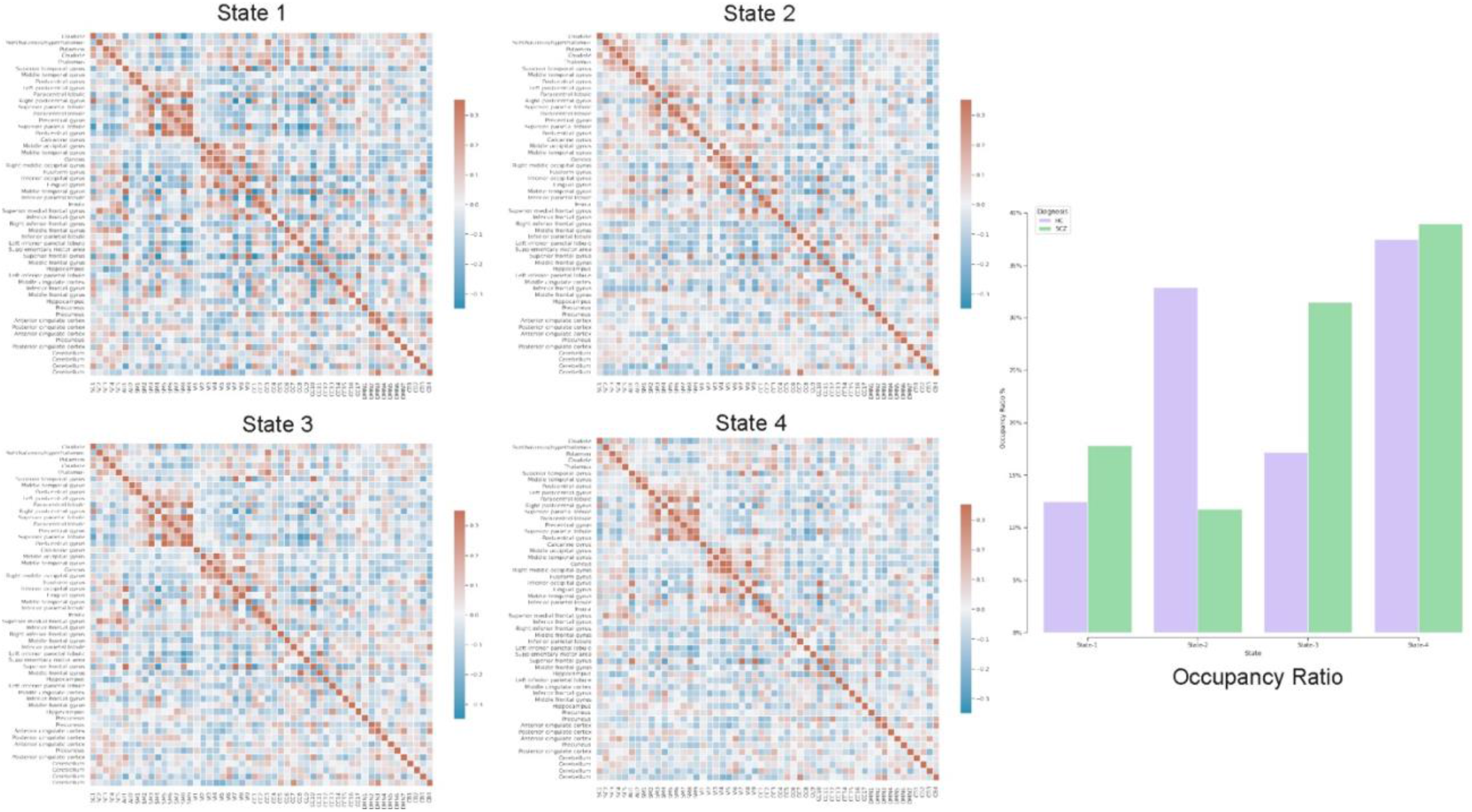
Each of states (Heatmaps) shows centroids, computed by k-means algorithm applied on all dFNC windows and the bar chart highlights temporal variation using occupancy ratio in health control versus schizophrenia subjects in 4 different states.

Our exploration of temporal brain dynamics illustrates high correlation between networks belonging to same domain while showing anti-correlation/weak correlation to others for both groups. We also see differences in the OR and behaviors of the dFNC maps. For example, we see different patterns in the centroids (4 heat maps) computed by k-means algorithm and control subjects spend more time in state-2 whereas schizophrenia patients spend more time in the other states.

## 4 Discussion

In this study, we conducted a group level analysis for characterizing 5D spatiotemporal networks in schizophrenia. We utilized the BPARC framework including 53 residual models to generate 4D score maps (probabilistic maps) for all subjects each of which is a representation of the relevant brain network. Our analysis demonstrates significant differences between control versus schizophrenia network representations in terms of space, time, and spatiotemporal brain dynamics.

### 4.1 Space, connectivity, and Spatiotemporal dynamism in schizophrenia

The proposed study, sought to extend findings supporting the validation of promising 4D representations generated by brain parcellation framework. We addressed the aforementioned goal by inspecting generated representation for schizophrenic patients in terms of spatial features, functional connectivity, and spatiotemporal brain dynamism.

Our observation shows significant differences in averaged spatial maps in which group level differences are lower in auditory and subcortical networks while sensori-motor, cerebellum, and default mode domains show the most differences with overall higher values in the patients. Moreover, it is also clear in figure 4 that we have higher median for peak voxels in positive segments the control group versus schizophrenia. Conversely, we find a higher median in negative segments for the schizophrenia group, suggesting overall higher activity in the controls. The PDF (probability density function) of the coefficients for both groups shows positive kurtosis in the negative segment of the controls for the cognitive control-1 network. Our results are in line with previous studies in schizophrenia reporting higher amplitude for default mode network in controls [26] or significant voxel-wise differences (higher t-value) for schizophrenic patients in subcortical and default mode network [27]. Moreover, we observed different patterns of spatial brain dynamism between controls versus schizophrenia subjects. Our result illustrates a lower level of voxel-wise variation over time for thalamus, hypothalamus, cerebellum, precuneus, anterior cingulate cortex, paracentral lobule, and frontal gyrus in schizophrenic patients. In another word, we can detect a partially semi-stationary status of spatial dynamics in cerebellum, default mode, subcortical, and cognitive control domains for schizophrenic patients in comparison with control subjects.

On the other side, plenty of research have reported an aberrant pattern in functional connectivity being associated with schizophrenia [3, 9]. According to our functional network connectivity measurements, we can see hyperconnectivity of different brain domains in schizophrenic patients which are consistent with recent findings like visual domain and subcortical domain [13], default mode with sensori-motor and subcortical domains [28]. Previous studies also reported hypoconnectivity of thalamus with frontal lobe [9, 29]. Thus, we detected thalamic hypoconnectivity with inferior frontal gyrus, superior frontal gyrus, and hippocampus which are similarly reported in previous studies as well. Another group of studies recognized transient reduction in temporal brain dynamism [15, 30] which is compatible with our results regarding dynamic FNC and dwell time of different states. Our observation supports the hypothesis of altered connectivity patterns in schizophrenia and possibility of these patterns with psychological interventions.

### 4.2 Advantages of the 4D maps

The proposed approach generates full 4D representations from input fMRI data and enables us to directly study spatiotemporal brain dynamism. A good feature of the proposed method is generating individual maps for each subject which makes it a better choice rather than a group of schizophrenia studies in which atlas-based parcellation plays a key role. Atlas-based analysis are widely used for studying potential biomarkers in schizophrenia by using a fixed size template for all subjects and then conducting statistical analysis on different segments [31, 32, 33], but obviously they ignore natural differences between brains in shape, size, and folding, which are considered in 4D approach.

A major advantage of the 4D network approach is we can capture changes which are only visible in the voxel level spatial dynamics. One dramatic example which highlights this feature is our observation of functional activity in the sensori-motor3 network that shows transient linking to the cerebellum as is shown in figure 8.

**Figure 8.**
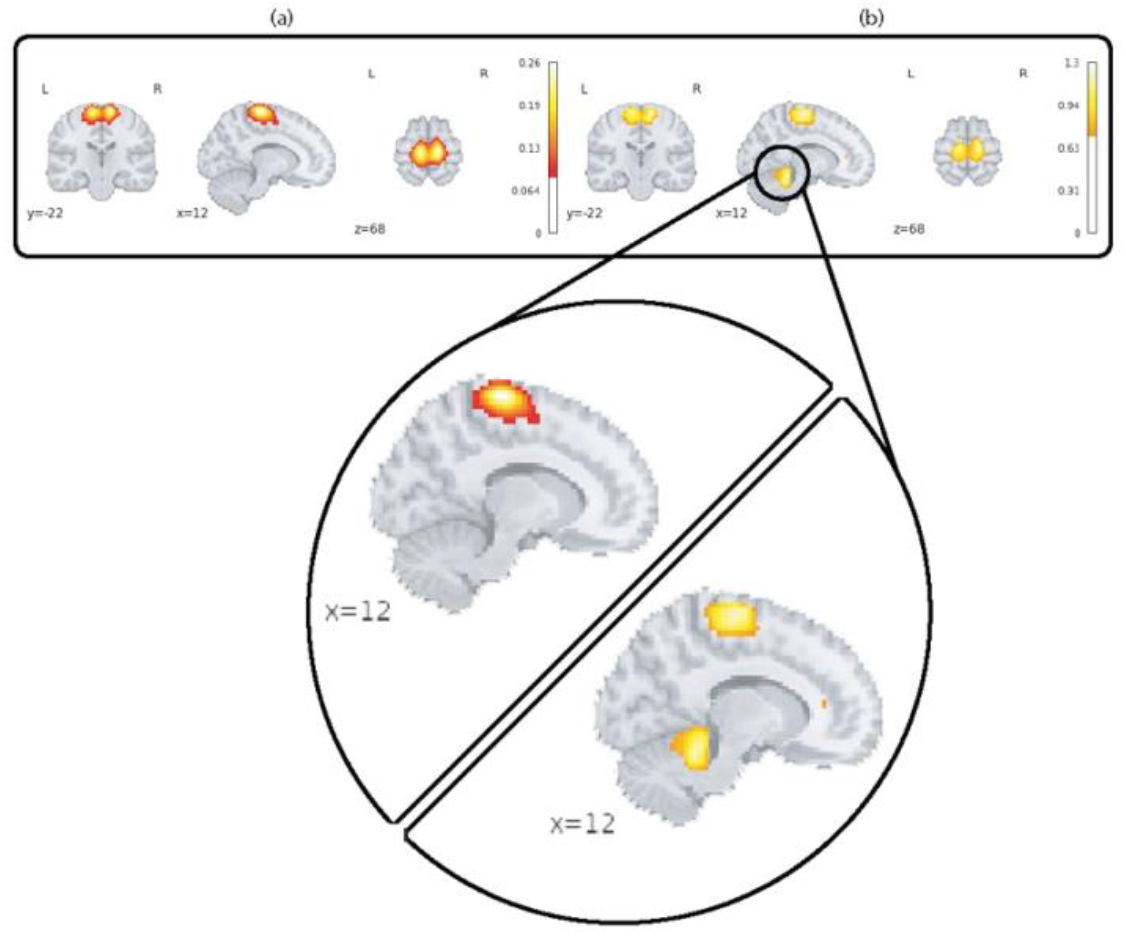
This figure highlights spatial dynamics over time in a sensori-motor network (SM3) by showing a transient state from sensori-motor to cerebellar network. Part (a) shows the averaged map over time and subjects for the control group and section (b) shows spatial deviation over time which is averaged over subjects for the same network/group.

This feature can open up a wide range of possibilities in studying brain disease and makes the 4D approach different from popular ICA-based analysis. In ICA-based methods, we can compute individual spatial components and time-courses for each subject, then applying statistical analysis on them [13, 34, 35], but variation of voxels in shape, size, and translation over time —spatial dynamics is a missing piece of the puzzle. Moreover, the proposed 4D approach has less computational complexity in inference phase comparing with ICA-based studies which can also consider a minor advantage as well.

### 4.3 Limitations

We can numerate several limitations and assumptions in the proposed schizophrenia study which might be considered in future research. However, we are able to relatively replicate plausible representation (score map) patterns for all subjects, it is not still clear whether observed patterns are generated by neuronal and cognitive sources originally or just that of artifacts which is a common challenge in recent research [9, 35]. It is not also feasible to evaluate impact of short and long-term pharmacologic treatment on changing neuronal activity patterns which are represented by the models for schizophrenia patients [36, 37]. Moreover, schizophrenia is a complex brain disease where different internal or environmental factors interact with each other to affect brain and eventually appears into clinical symptoms which not investigated in our study including gender, age, genetics, IQ, etc [38, 39].

## 5 Conclusion

In this work, we have studied group level differences in spatiotemporal brain dynamics between healthy-control and schizophrenia subjects by incorporating BPARC framework including 53 different pre-trained models. Our testing and evaluations show group level differences across multiple networks and spatiotemporal features which need further study as a potential brain-based biomarker for schizophrenia. In future, we will utilize generated representations for classifying control and schizophrenia subjects and studying impact and contribution of different brain networks in schizophrenia. Also, we will incorporate other factors like gender, age, and ethnicity in our analysis including during the model training process, to more fully address potentially confounding factors.

## 7 Author Contributions

**Behnam Kazemivash:** Methodology, Software, Investigation, Writing. **Theo GM VanErp:** Resources, Writing. **Peter Kochunov:** Resources, Writing. **Vince D. Calhoun:** Supervision, Conceptualization, Resources, Writing.

## 9 Acknowledgements

The BPARC framework along with all analysis and implementation are available under a MIT license, via https://github.com/bkazemivash/BPARC.

